# Brain network organization and postoperative cognitive change in meningioma and glioma

**DOI:** 10.64898/2026.05.17.725702

**Authors:** Albert Juncà, Ignacio Martín, Gustavo Deco, Gustavo Patow

## Abstract

Brain tumours can alter connectivity at both regional and whole-brain scales. We examined how brain networks fitted with a whole-brain model differ across tumour groups, and whether postoperative changes in functional network organization accompany cognitive change. We performed an exploratory secondary analysis of structural and resting-state functional MRI and cognitive data from 14 participants with meningioma, 11 in the glioma group and 11 controls. Paired preoperative and follow-up assessments were available for 27 participants. Generative effective connectivity was estimated by fitting the model to two functional targets, functional connectivity and time-lagged covariance. Fitted networks in the glioma group showed lower weighted clustering than those in controls and meningiomas (adjusted *P* = 0.021 and *P* = 0.026, respectively). Their fitted networks also had higher incoming participation, indicating a more even distribution of incoming connection strength across network communities. In sensitivity analyses, the difference persisted when each participant’s own structural connectivity was used for fitting, and mean clustering was similar across groups when all participants were fitted to the same control-average functional targets. In all ten eligible glioma participants, average functional connectivity between tumour-overlapping regions and their contralateral counterparts was below the control mean, both without and with global signal regression. Among ten meningioma participants with paired assessments, changes in functional connectivity modularity, which measures how strongly a network divides into communities, showed positive but inconclusive correlations with overall cognitive change. An exploratory analysis of two cognitive subscores showed that modularity change without global signal regression was positively associated with change in the attention/ planning/spatial-span score (Spearman’s *ρ* = 0.79, adjusted *P* = 0.026). This association remained after accounting for preoperative cognition and modularity. With global signal regression, the corresponding association was weaker and inconclusive. These findings characterize differences in fitted network organization across tumour groups. Regional connectivity provides anatomical context in gliomas, while longitudinal measurements link changes in functional connectivity modularity to cognitive change in meningiomas. These observations need confirmation in larger, independent cohorts.

## 1 Introduction

Brain tumours can affect connectivity through several processes, including infiltration and compression of surrounding tissue. Diffuse gliomas infiltrate the brain parenchyma [1]. Meningiomas arise from the meninges and usually grow extra-axially, compressing adjacent brain tissue [2]. Brain invasion can nevertheless occur in meningiomas [2], and peritumoral oedema has been associated with altered functional connectivity (FC) in these patients [3]. These anatomical differences provide a rationale for comparing network organization across tumour groups.

Connectivity changes are not necessarily confined to the lesion. In glioma, altered resting-state network topography has been reported beyond MRI-defined tumour and oedema boundaries [4]. Prior work using the dataset analysed here reported higher structural participation coefficients in gliomas than in controls, while group differences in structural global efficiency and modularity were not statistically significant [5]. Another analysis of these data examined intrinsic ignition and metastability, identifying group differences in functional dynamics [6]. Earlier longitudinal work in this cohort examined the stability of fitted model parameters after resection [7]. A complementary question is how regional connectivity differences relate to whole-brain network organization.

Structural connectivity (SC) constrains but does not fully determine FC: regions without a detected direct structural link can still be functionally coupled [8, 9]. Whole-brain models explore this relationship by describing how regional dynamics interact through anatomical connections [5]. The model’s coupling weights can then be fitted to selected functional observations, yielding estimates of generative effective connectivity (GEC) [10]. We characterize the fitted GEC networks using clustering and participation, and check whether group differences hold under different structural and functional inputs. Clustering describes interconnections among a region’s network neighbours, while participation describes how its connection strength is distributed across communities [11]. Regional SC and FC measurements provide anatomical context for these network comparisons.

Longitudinal measurements also allow changes in network organization to be related to cognitive change. FC modularity quantifies within-community connectivity relative to a network null model [12]. In a heterogeneous surgical cohort assessed before and approximately two weeks after resection, Ellis et al. reported a positive association between changes in FC modularity and changes in a neuropsychological composite score in a multivariable analysis [13]. Their finding motivates examining FC–cognition associations at later follow-up and within tumour groups. We also examine whether modularity–cognition associations are retained in model-generated FC. Because measured FC enters model fitting, this comparison serves as an exploratory consistency check.

Here, we perform an exploratory secondary analysis of preoperative and follow-up MRI and cognitive data from meningioma and glioma participants and controls (Fig. 1). First, we compare the organization of GEC networks across groups and examine its sensitivity to structural and functional model inputs (Fig. 1B). Second, in eligible glioma participants, we compare SC strength between tumour-overlapping regions and their contralateral counterparts, measure FC between the paired regions, and relate regional SC differences to global GEC clustering (Fig. 1C). Third, we examine whether changes in FC modularity accompany cognitive change at follow-up, with detailed analyses in meningiomas (Fig. 1D). Because the global fMRI signal contains neural and non-neural contributions, we examine empirical FC with and without global signal regression (GSR) as complementary views of connectivity [14].

**Figure 1.**
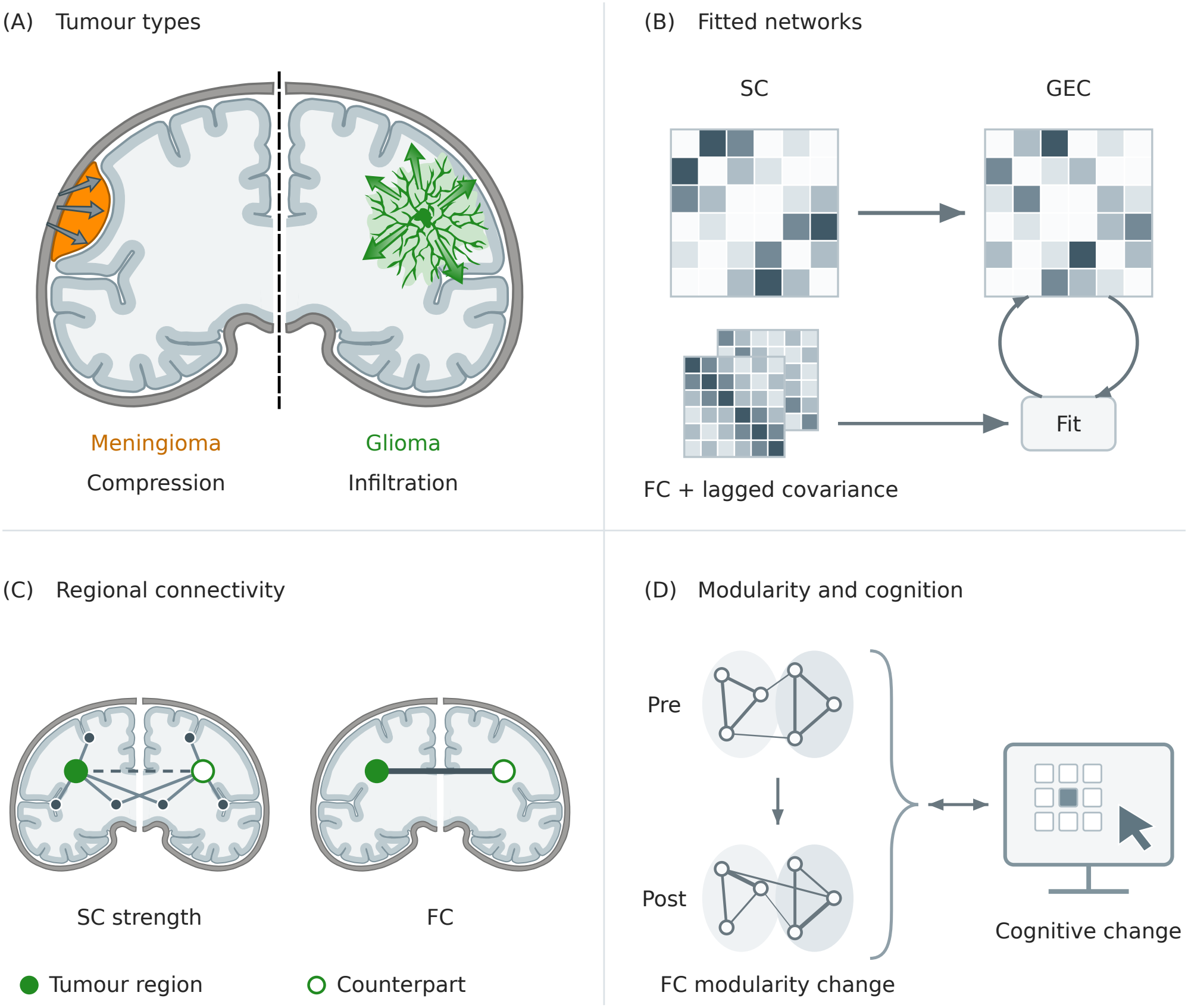
Overview of the study. **(A)** Schematic anatomical contrast between meningioma compression and glioma infiltration. **(B)** Common control SC provides the starting matrix. Fitting iteratively adjusts GEC weights to match model FC and normalized time-lagged covariance to their empirical counterparts. Both functional targets are calculated directly from BOLD signals. **(C)** In glioma participants, we compare SC strength between tumour regions and their counterparts in the opposite hemisphere. We measure FC between the two regions in each pair. We also assess whether the SC strength difference is associated with global GEC clustering. **(D)** We assess whether changes in FC modularity are associated with cognitive changes, separately within each group. Further analyses focus on meningiomas. All illustrations are schematic. SC, structural connectivity; FC, functional connectivity; GEC, generative effective connectivity.

## 2. Materials and methods

### 2.1 Study design and participants

We performed an exploratory secondary analysis of the publicly available BTC_preop and BTC_postop datasets, combining preoperative group comparisons with longitudinal analyses. We used OpenNeuro datasets ds001226, version 5.0.1, and ds002080, version 4.0.1 [15, 16]. Sample size was determined by data availability.

Participants were recruited at Ghent University Hospital, Belgium, between May 2015 and October 2017. Original patient eligibility criteria included age *≥*18 years, a supratentorial meningioma (WHO grade I–II) or glioma (WHO grade II–III), ability to complete neuropsychological testing, and medical clearance for MRI. Controls were recruited among patients’ caregivers. Assessments were performed before surgery and at follow-up several months later, with repeated assessments in controls. The original study received approval from the Ethics Committee at Ghent University Hospital, and all participants provided written informed consent [17, 7].

The preoperative connectivity sample comprised 36 participants: 14 with meningioma, 11 in the glioma group and 11 controls. We retained the dataset’s diagnostic grouping; one participant in the glioma group was recorded as having a grade II ependymoma. Demographic and clinical characteristics, together with cognitive data availability, are reported in Table 1.

**Table 1.** Participant characteristics and follow-up availability.

| Characteristic | Controls | Meningiomas | Glioma |
|---|---|---|---|
| Preoperative participants, *n* | 11 | 14 | 11 |
| Age (years), mean (SD) | 58.6 (10.3) | 60.4 (12.3) | 47.5 (11.3) |
| Female, *n* (%) | 4 (36.4) | 11 (78.6) | 4 (36.4) |
| Tumour volume (cm<sup>3</sup>), median [Q1, Q3] | — | 3.0 [1.8, 15.8] | 15.2 [13.1, 35.1] |
| Preoperative cognitive composite, *n* | 11 | 14 | 10 |
| Postoperative FC and cognition, *n* | 10 | 10 | 7 |

SD, sample standard deviation; Q1 and Q3, first and third quartiles; FC, functional connectivity. Demographic and tumour-volume summaries refer to the preoperative cohort. Percentages use the full preoperative group size; a dash denotes not applicable. Postoperative counts include participants with data at both assessments; controls underwent repeat assessment. Glioma includes one participant recorded as grade II ependymoma. One meningioma participant contributes six of seven cognitive outcomes.

Follow-up records were available for 29 participants. Two lacked the supplied postoperative functional-connectivity derivative, leaving 27 participants with paired functional connectivity (FC) and cognitive measurements: 10 meningiomas, seven gliomas and 10 controls. Postoperative FC–cognition associations were assessed separately within each group. The main longitudinal presentation focuses on meningiomas, including the additional baseline-adjusted analyses.

### 2.2 MRI data and preprocessing

MRI data were acquired on a 3 T Siemens Magnetom Trio scanner with a 32-channel head coil. Acquisitions included T1-weighted anatomical imaging, multi-shell diffusion imaging and resting-state functional MRI, with isotropic voxel sizes of 1, 2.5 and 3 mm, respectively. The repetition time (TR) of the resting-state scans was 2.1 or 2.4 s, because the protocol changed during recruitment. Full acquisition details are provided in the dataset description [17].

We used two derivatives supplied with the dataset: structural connectivity matrices and regional blood-oxygen-level-dependent (BOLD) time series. Both follow the Desikan–Killiany parcellation of 68 cortical regions, 34 per hemisphere [18]. The dataset authors created each participant’s parcellation with FreeSurfer. In meningioma participants, lesions were filled where required. In glioma participants, the postoperative parcellation was obtained by registering the preoperative parcellation to the postoperative anatomy [7].

Structural connectivity was reconstructed by the dataset authors using multi-tissue constrained spherical deconvolution, anatomically constrained probabilistic tractography and spherical-deconvolution informed filtering of tractograms. Thirty million streamlines were generated per participant and reduced to 7.5 million. Connection weights were streamline counts between regions. Connections with fewer than five streamlines were removed, and all weights were divided by 75,000 [17]. We used these matrices as supplied; rescaling for model fitting is described in Section 2.3.

The dataset authors preprocessed the functional images with FSL FEAT, including motion correction, slice-timing correction, brain extraction, intensity normalization and high-pass filtering with a 100 s cutoff. They then mapped each cortical parcellation to functional space to extract the regional BOLD signals [7].

Our own processing started from these regional signals. We removed the linear trend and mean of each signal, then applied a Butterworth band-pass filter of 0.01–0.08 Hz (SciPy order parameter *N* = 2). The filter was run forward and backward, using each scan’s own TR. We applied the same steps at both assessments and kept all 180 supplied time points.

FC was the Pearson correlation between each pair of regional signals; negative correlations were retained. We calculated it twice, without and with global signal regression (GSR). Without GSR, FC was calculated directly from the filtered signals. For GSR, we first regressed the global signal and an intercept from each regional signal, with the global signal defined as the equal-weight mean of the 68 filtered signals. No further filtering followed the regression. The two are complementary descriptions of FC; reporting both shows how far a result depends on signal shared across the cortex [14].

### 2.3 Generative effective connectivity

Generative effective connectivity (GEC) was estimated using a linearized Hopf whole-brain model following Deco et al. [19]. Each cortical region was represented by a coupled oscillator, with its real-valued activity representing BOLD fluctuations. The bifurcation parameter was fixed at *a* = *−*0.02, placing the local dynamics below the onset of self-sustained oscillations. Independent Gaussian noise had amplitude *σ* = 0.01. Model FC and time-lagged covariance were calculated analytically from the linearized dynamics.

For the primary analysis, all participants shared one structural starting matrix, the common control SC. It was the mean SC of the 11 controls, rescaled to a maximum weight of 0.3, with diagonal entries and weights below 10^*−*6^ set to zero. Regional intrinsic frequencies were also shared. For each region, the peak frequency of each control’s smoothed BOLD power spectrum was averaged within each TR group, and the two group means were then averaged with equal weight.

Each participant’s GEC matrix was fitted to two functional targets calculated from their filtered BOLD signals without GSR: FC and normalized time-lagged covariance. The lag was two sampling intervals (4.2 or 4.8 s, depending on TR). Lagged covariances were normalized by the product of the two regions’ standard deviations at zero lag. Connection weights were adjusted using the pseudo-gradient update

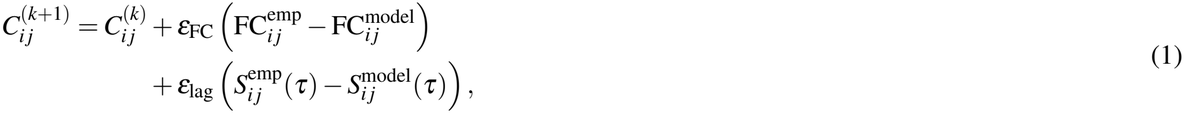

where *S*(*τ*) denotes normalized time-lagged covariance and *C*_*i j*_ represents coupling from region *j* to region *i*. The update coefficients were *ε*_FC_ = 0.0004375 and *ε*_lag_ = 0.0000625, giving a ratio of 7:1.

After each update, negative weights were set to zero and the matrix was rescaled to a maximum weight of 0.3. Connections at zero stayed at zero. Fitting ran for exactly 10,000 updates per participant, and the final matrix was retained. Fitting settings were selected during exploration of this cohort and applied uniformly to all participants. Fit quality was measured as the mean squared error of FC and of normalized time-lagged covariance. All fitted models were stable, with every Jacobian eigenvalue having a negative real part. For the longitudinal analysis, GEC was fitted separately at each assessment, using the same preoperative control starting matrix, frequencies and fitting settings. As sensitivity analyses, we repeated the fitting with each participant’s own SC as the starting matrix, rescaled and thresholded in the same way, with all other settings unchanged. Each of these matrices was fitted twice: to the participant’s own functional targets and to control-average targets. Control-average FC was the mean of the 11 control FC matrices. Control-average normalized lagged covariance was calculated within each TR group, and each participant was fitted to the average for their TR. For each participant, we then subtracted global GEC clustering with control-average targets from clustering with their own targets.

### 2.4 Network measures

Each GEC matrix was analysed as a weighted, directed network of all 68 cortical regions, keeping every positive connection without further thresholding. Clustering followed the weighted, directed definition of Fagiolo [20]: it measures how many of the possible triangles around each region are present, with each triangle weighted by the geometric mean of its connection weights. For clustering, weights were first divided by the largest weight in the network, and the 68 regional coefficients were averaged. Global efficiency, a measure of connectivity through weighted paths, was calculated from the fitted weights as the mean inverse shortest-path length over all ordered pairs of distinct regions [11]. The length of a connection was the reciprocal of its weight, and pairs with no directed path contributed zero.

The participation coefficient measures how evenly a region’s connection strength is spread across network communities; higher values indicate a more even spread [11]. Communities were identified separately for each GEC network using NetworkX’s directed, weighted implementation of the Louvain algorithm with resolution parameter *γ* = 1 [21]. Participation was calculated as

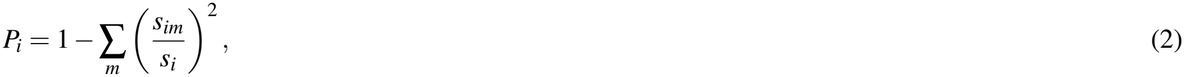

where *s*_*im*_ is the connection strength between region *i* and community *m*, and *s*_*i*_ is the region’s total strength. Incoming and outgoing participation were calculated separately from incoming and outgoing strengths, and regions with zero strength in that direction were assigned zero. Regional coefficients were averaged across the 68 regions. Because Louvain results vary between runs, this network average was itself averaged across 20 runs with fixed random seeds (0–19).

The longitudinal analysis used FC modularity and was motivated by Ellis et al. [13]. Each FC matrix was converted to an undirected, weighted network containing all 68 regions and the 456 strongest positive correlations among the 2,278 possible connections. This density of approximately 20% followed their study. Retained connections kept their Pearson correlations as weights, and all other entries were set to zero. Their study used predefined Yeo seven-network communities; we instead identified communities separately for each network, using the Louvain algorithm with *γ* = 1. Modularity compares connection strength within communities with that expected from the regions’ total strengths [21]. It was calculated for each participant at each assessment, with and without GSR, and averaged across 20 Louvain runs with the same seeds. Longitudinal change was postoperative minus preoperative modularity. The same modularity calculation was applied to model-generated FC, the FC produced by the fitted GEC networks.

Clustering, community detection and modularity were calculated with NetworkX 3.6.1 [22].

### 2.5 Regional connectivity analyses

Regional tumour involvement was estimated from the dataset authors’ MNI-space tumour masks and a common Desikan–Killiany atlas derived from FreeSurfer 7.4.1’s fsaverage parcellation. The atlas was transformed to the FSL MNI152 1 mm grid using FreeSurfer’s supplied MNI152 registration. Tumour masks were binarized at values greater than 0.5 and resampled to the same grid. Nearest-neighbour interpolation was used throughout. The tumour overlap of a region was the fraction of that atlas region occupied by the tumour mask.

We selected tumour regions, defined as regions with more than 10% tumour overlap whose counterpart, the same region in the opposite hemisphere, had zero overlap. This gave 33 tumour–counterpart pairs in ten glioma-group participants; one participant had no eligible region. Regional analyses focused on gliomas because only four meningioma participants met these criteria. The counterpart served as an anatomical comparator and was not assumed to be unaffected. As a sensitivity analysis, SC results were also calculated without the zero-overlap requirement for the counterpart.

Regional SC strength was defined as the mean connection weight between each region and the other 67 cortical regions. Strengths were standardized as *z* = (*x− µ*_C_)*/σ*_C_, where *µ*_C_ and *σ*_C_ are that region’s mean and population standard deviation across the 11 controls. For each selected pair, we subtracted the counterpart’s SC *z*-score from the tumour region’s SC *z*-score, then averaged these differences within each patient. Negative values indicate lower standardized SC strength at tumour regions than at their counterparts. We used Spearman correlation to assess whether these SC strength differences were associated with global GEC clustering (Section 2.4) in the networks fitted from the common control SC.

For regional FC, we took the FC between each tumour region and its counterpart, without and with GSR. Each value was standardized against the same connection in the 11 controls, using the *z*-score above within the same GSR condition. These *z*-scores were averaged within each patient; negative values indicate lower FC than the control mean. SC and FC used the same tumour–counterpart pairs, and each patient contributed equally to the group summaries.

We also explored whether SC strength differences varied with tumour overlap. This analysis used every region– counterpart pair with any tumour overlap, 68 patient-specific pairs in 11 glioma-group participants, taking the side with greater overlap as the tumour side. We related each pair’s SC strength difference to its difference in tumour overlap, using one common slope after centring both variables within each patient. One participant had a single pair and therefore did not contribute to the slope.

### 2.6 Cognitive assessment

Cognitive performance was assessed using the Cambridge Neuropsychological Test Automated Battery (CANTAB), as described in the dataset publication [17]. We used seven outcome measures from five tasks: mean latency from Motor Screening (MOT); simple movement time, simple reaction time and five-choice reaction time from Reaction Time (RTI); sustained-attention sensitivity (*A*^*′*^) from Rapid Visual Information Processing (RVP); problems solved in the minimum number of moves from Stockings of Cambridge (SOC); and span length from Spatial Span (SSP).

Each outcome was standardized using the mean and sample standard deviation of the 11 controls at baseline. The four timing measures from MOT and RTI were sign-reversed, so that higher scores always indicated better performance. The overall cognitive composite was the unweighted mean of the available standardized outcomes.

For longitudinal analyses, scores at both assessments were standardized against the same baseline controls, and each participant’s scores used only the outcomes available at both assessments. One meningioma participant lacked baseline RVP data and therefore contributed six outcomes at each assessment; the remaining paired participants contributed all seven. Missing values were not imputed. Cognitive change was the postoperative score minus the preoperative score.

For an exploratory follow-up of the modularity–cognition association, we also calculated two subscores. The latency score averaged the four MOT and RTI outcomes, and the attention/planning/spatial-span score averaged the RVP, SOC and SSP outcomes. The composite and both subscores are exploratory summaries and have not been validated as clinical scales.

### 2.7 Statistical analysis

The participant was the unit of analysis. The regional SC and FC summaries and the SC–GEC association used one average value per patient. All tests were two-sided. Confidence intervals were 95% and were not adjusted for multiple comparisons. Network measures were summarized as means and standard deviations. *P*-values were adjusted with the Benjamini–Hochberg (BH) procedure within each set of related tests, and reported *P*-values are BH-adjusted unless stated otherwise. Values below 0.05 were considered statistically significant. Table 2 lists each analysis with its participants, test, interval method and correction set. Analyses used Python, SciPy 1.15.0 [23] and statsmodels 0.14.4.

**Table 2.**
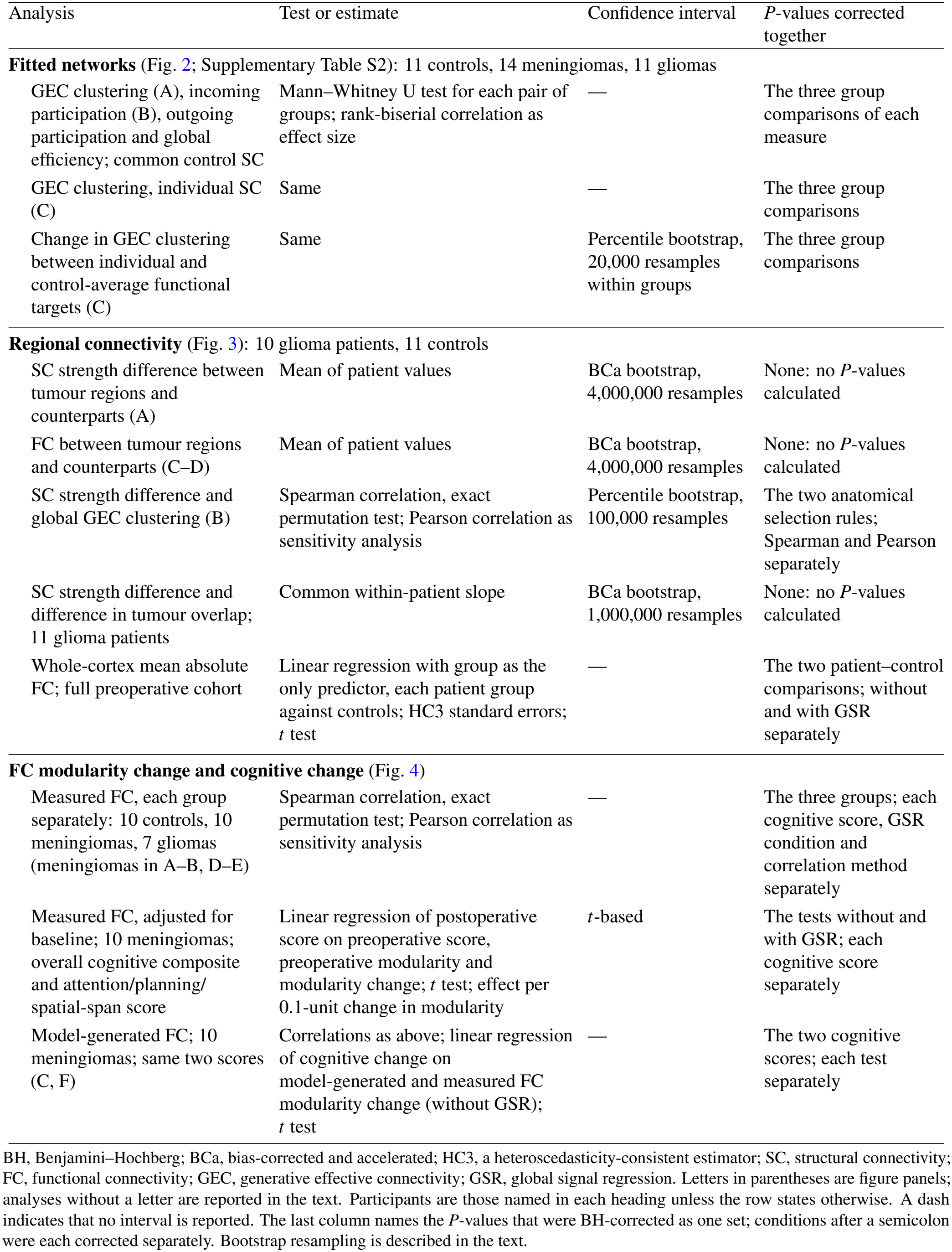
Statistical analyses.

Bootstrap intervals resampled whole participants. For the regional analyses, patients and controls were resampled independently and the control means and standard deviations were recalculated in every resample, so these intervals include uncertainty in the control reference. The exact permutation test for the SC–GEC association held that reference fixed, so its *P*-value and its interval do not come from the same procedure. The number of resamples (Table 2) was set separately for each analysis. Bias-corrected and accelerated (BCa) limits fall far in the tails of the bootstrap distribution and needed more resamples to be numerically stable.

Primary group comparisons did not adjust for age, sex or acquisition TR. For whole-cortex FC and the GEC functional-target difference, we also assessed sensitivity to adjustment for age and TR, and additionally for sex (Supplementary Methods and Table S1).

The BH corrections apply to the sets listed in Table 2, not to all analyses in the paper or to the exploration that preceded them. Analyses with and without GSR used the same participants and are not independent replications.

## 3 Results

Preoperative analyses included 11 controls, 14 participants with meningioma and 11 participants in the glioma group. Longitudinal FC–cognition analyses included 27 participants with paired assessments: 10 controls, 10 with meningioma and seven in the glioma group (Table 1).

### 3.1 Fitted network organization across tumour groups

We examined whether the organization of fitted GEC networks differed across controls and the two tumour groups (Supplementary Table S2). Each network was fitted from the common control SC matrix to that participant’s functional targets. Weighted clustering was lower in the glioma group than in controls and meningiomas (Fig. 2A). Clustering was 0.108 *±* 0.019 in gliomas, 0.145 *±* 0.027 in controls and 0.131 *±* 0.026 in meningiomas (mean *±* SD). The glioma–control and meningioma–glioma comparisons gave *P* = 0.021 and *P* = 0.026, respectively. The meningioma–control comparison was inconclusive (*P* = 0.239).

**Figure 2.**
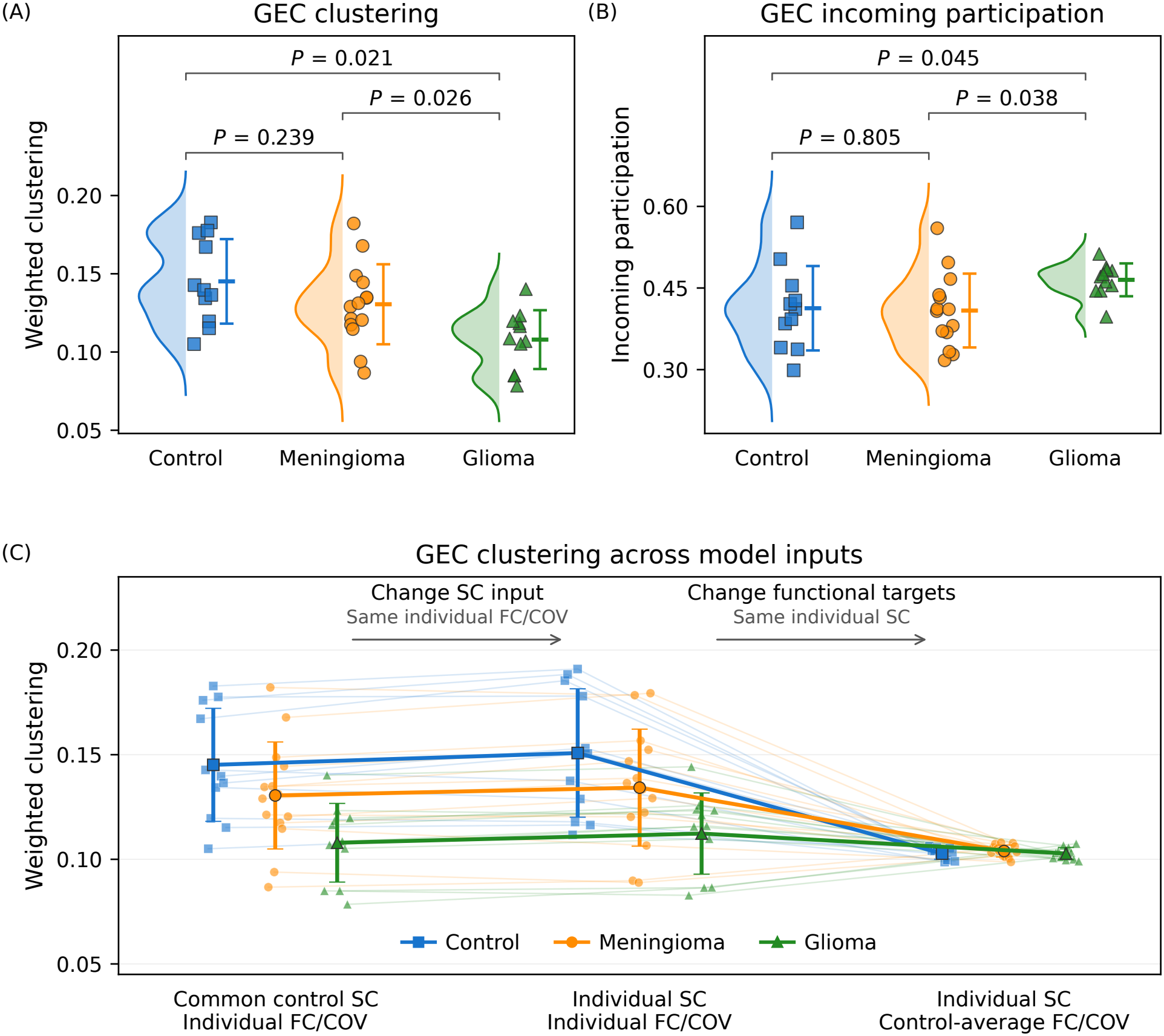
Fitted GEC network organization and sensitivity to model inputs. **(A)** Weighted clustering and **(B)** incoming participation in networks fitted using common control SC and individual functional targets. Symbols represent participants: controls (blue; *n* = 11), meningiomas (orange; *n* = 14) and gliomas (green; *n* = 11). Half-violins show distributions; bars show means *±*SD. Brackets show BH-adjusted *P*-values across the three group comparisons within each measure. **(C)** Clustering in the same participants under three preoperative fitting conditions: common control SC with individual functional targets; individual SC with individual functional targets; and individual SC with control-average functional targets. FC/COV denotes the functional targets used for fitting: FC and normalized time-lagged covariance. Thin lines connect each participant’s values; larger symbols and thick lines show group means with SD bars. Arrows identify the input changed between conditions. Panel C describes fitting conditions, not longitudinal change. All networks were fitted without GSR.

Incoming participation was higher in gliomas than in controls and meningiomas (Fig. 2B). Participation was 0.465 *±*0.030 in gliomas, 0.413*±* 0.078 in controls and 0.409 *±*0.068 in meningiomas. The glioma–control and meningioma–glioma comparisons gave *P* = 0.045 and *P* = 0.038, respectively. The meningioma–control comparison was inconclusive (*P* = 0.805).

Outgoing participation was also higher on average in gliomas than in controls and meningiomas, although the glioma–control and meningioma–glioma comparisons were inconclusive (*P* = 0.063 and *P* = 0.060, respectively). Global-efficiency comparisons were also inconclusive: glioma–control, *P* = 0.065; meningioma–control, *P* = 0.088; and meningioma–glioma, *P* = 0.262.

When each participant’s own SC replaced the common control SC as the starting matrix, with the same functional targets, clustering remained lower in gliomas than in controls (*P* = 0.021) and meningiomas (*P* = 0.046; Fig. 2C). The meningioma–control comparison remained inconclusive (*P* = 0.286).

With each participant’s SC held fixed, we then compared GEC clustering between fits to individual and control-average functional targets (Fig. 2C). Mean clustering values were closer across groups with control-average functional targets: 0.103 in controls, 0.104 in meningiomas and 0.103 in gliomas. For each participant, we calculated clustering with individual functional targets minus clustering with control-average functional targets. This within-participant clustering difference averaged 0.048 in controls, 0.030 in meningiomas and 0.010 in gliomas. The clustering difference was smaller in gliomas than in controls (*P* = 0.026) and meningiomas (*P* = 0.040). The meningioma–control comparison was inconclusive (*P* = 0.163).

### 3.2 Regional connectivity and its association with fitted-network organization

To provide anatomical context for the fitted-network findings, we examined SC and FC in 33 tumour–counterpart pairs from ten glioma-group participants (Fig. 3). We compared SC strength between the two regions of each pair and measured the FC between them.

**Figure 3.**
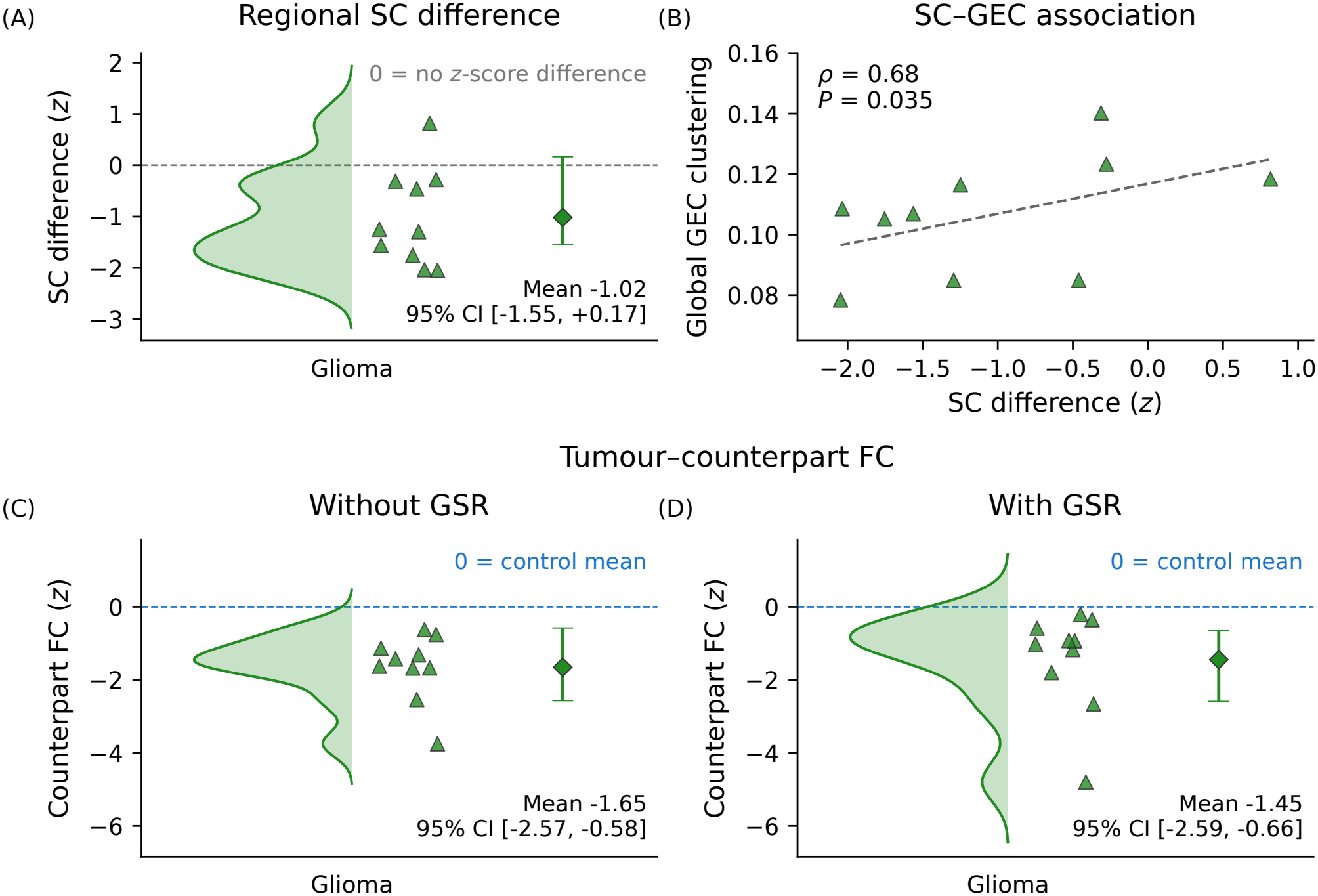
Regional connectivity and its association with global GEC clustering. All panels use the same 33 tumour–counterpart pairs from ten glioma-group participants, selected as described in the Methods. Each point represents one participant. **(A)** Control-standardized SC strength differences between tumour regions and their counterparts, averaged within each patient. Negative values indicate lower standardized strength at tumour regions. **(B)** Association between the SC strength differences in A and global GEC clustering. Spearman’s *ρ* and the BH-adjusted *P*-value are shown. The dashed line is a linear fit shown as a visual guide. **(C–D)** FC values for tumour–counterpart connections, standardized against the same connections in controls, without GSR (C) and with GSR (D). Zero indicates the control mean. In A, C and D, half-violins show participant distributions; diamonds and error bars show means and 95% confidence intervals.

In nine of the ten patients, standardized SC strength was on average lower at tumour regions than at their counterparts. The mean SC strength difference was *−*1.02 *z*-score units (95% CI, *−*1.55 to 0.17; Fig. 3A), although the interval included zero. As a sensitivity analysis, we removed the zero-overlap requirement for the counterpart, which gave 35 pairs in the same ten patients. The mean SC strength difference was then *−*1.07 *z*-score units (95% CI, *−*1.55 to *−*0.16).

We also examined whether greater differences in tumour overlap between the two regions of a pair were associated with more negative SC strength differences. This exploratory analysis used every region–counterpart pair with any tumour overlap: 68 patient-specific pairs in 11 glioma-group participants. The estimated within-patient slope was *−*0.18 *z*-score units per additional 10 percentage points of overlap difference (95% CI, *−*0.31 to 0.07), although the interval included zero.

Among the ten patients in the main selection of 33 pairs, those with more negative SC strength differences tended to have lower global GEC clustering in the networks fitted from the common control SC (Spearman’s *ρ* = 0.68, *P* = 0.035; Fig. 3B). The bootstrap interval was wide and included zero (95% CI, *−*0.18 to 0.93). The Pearson correlation was also positive but inconclusive (*r* = 0.48, *P* = 0.165). The broader selection of 35 pairs gave the same Spearman correlation.

In all ten patients, FC between tumour regions and their counterparts was on average lower than the control mean, both without and with GSR (Fig. 3C–D). The group mean FC *z*-score was *−*1.65 without GSR (95% CI, *−*2.57 to *−*0.58) and *−*1.45 with GSR (95% CI, *−*2.59 to *−*0.66).

For context, we also compared whole-cortex mean absolute FC, the mean absolute correlation over all 2,278 region pairs, between each patient group and controls in the full preoperative cohort. It was lower in gliomas than in controls with GSR (difference *−*0.062, *P* = 0.007); without GSR the difference was similar in size but inconclusive (*−*0.068, *P* = 0.311). The meningioma–control comparisons were inconclusive in both conditions (*−*0.055 without GSR, *P* = 0.311; *−*0.010 with GSR, *P* = 0.683).

### 3.3 Postoperative changes in FC modularity and cognition

We examined, separately in each group, whether changes in FC modularity between the preoperative and postoperative assessments were associated with cognitive change. In the ten meningioma participants with paired assessments, correlations with the overall cognitive composite were positive both without GSR (Spearman’s *ρ* = 0.75, *P* = 0.052; Fig. 4A) and with GSR (*ρ* = 0.73, *P* = 0.061; Fig. 4B), but both were inconclusive after correction across the three groups. Pearson correlations showed a similar pattern (without GSR: *r* = 0.68, *P* = 0.088; with GSR: *r* = 0.68, *P* = 0.092).

**Figure 4.**
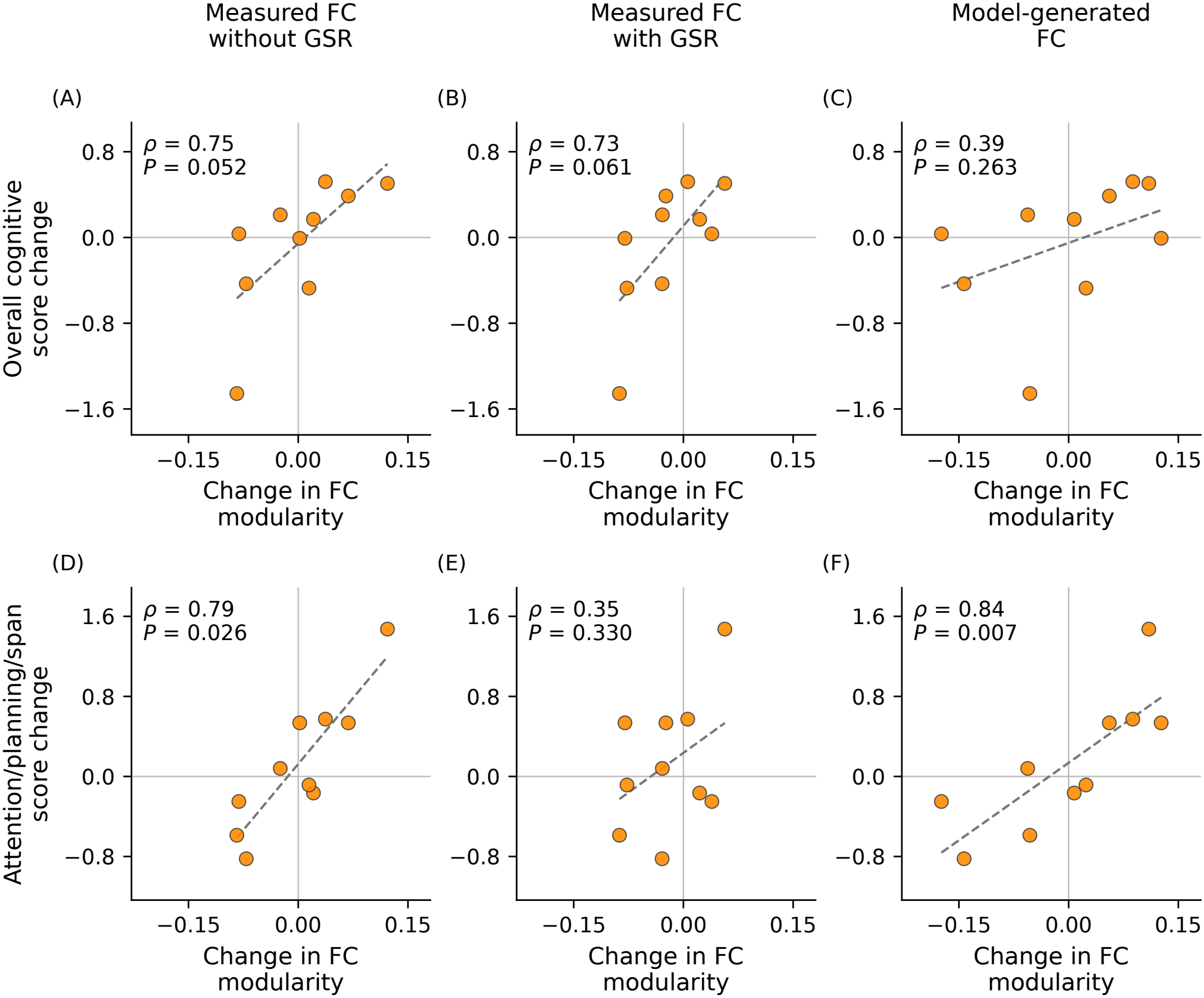
Changes in FC modularity and cognition after surgery. All panels show the same ten meningioma participants, with one point per participant. Changes are postoperative minus preoperative; higher cognitive scores indicate better performance. **(A–C)** Overall cognitive composite. **(D–F)** Attention/planning/spatial-span score. Columns show FC modularity change from measured FC without GSR (A, D), measured FC with GSR (B, E) and model-generated FC (C, F). In C and F, modularity is that of the model-generated FC, not of the GEC weights. Annotations show Spearman’s *ρ* and BH-adjusted *P*-values. BH correction covered the three groups for each measured-FC panel and the two cognitive scores for model-generated FC. Dashed lines are linear fits shown as visual guides; grey lines mark zero change.

To explore this pattern further, we assessed the latency score and the attention/planning/spatial-span score separately. In meningiomas, FC modularity change without GSR was positively associated with change in the attention/planning/spatial-span score (Spearman’s *ρ* = 0.79, *P* = 0.026; Fig. 4D). The corresponding association with GSR was inconclusive (*ρ* = 0.35, *P* = 0.330; Fig. 4E). Pearson correlations showed a similar pattern (without GSR: *r* = 0.89, *P* = 0.002; with GSR: *r* = 0.40, *P* = 0.250). Associations with the latency score were inconclusive both without GSR (*ρ* = 0.37, *P* = 0.492) and with GSR (*ρ* = 0.59, *P* = 0.242). Pearson correlations with the latency score were also inconclusive (without GSR: *r* = 0.34, *P* = 0.513; with GSR: *r* = 0.66, *P* = 0.111). In controls and glioma-group participants, the correlations with all three scores were inconclusive (Spearman and Pearson; smallest *P* = 0.111).

In meningiomas, we then tested whether these associations remained after accounting for the preoperative cognitive score and preoperative FC modularity. Without GSR, changes in modularity remained positively associated with postoperative attention/planning/spatial-span scores. The regression coefficient was 0.98 cognitive-score units per 0.1-unit change in modularity (95% CI, 0.54 to 1.43; *P* = 0.003). With GSR, the estimate was similar in magnitude but substantially less precise (0.92; 95% CI, *−*1.71 to 3.55; *P* = 0.426). For the overall cognitive composite, the coefficient was 0.69 without GSR (95% CI, 0.12 to 1.26) and 0.25 with GSR (95% CI, *−*1.47 to 1.97). Both associations were inconclusive after correction (*P* = 0.051 and *P* = 0.737).

As an exploratory consistency check, we repeated the analysis in meningiomas with model-generated FC from the networks fitted at each assessment. The association with the overall cognitive composite was inconclusive (Spearman’s *ρ* = 0.39, *P* = 0.263; Pearson’s *r* = 0.42, *P* = 0.233; Fig. 4C). Model-generated FC modularity change was positively associated with change in the attention/planning/spatial-span score (Spearman’s *ρ* = 0.84, *P* = 0.007; Fig. 4F), with a similar Pearson result (*r* = 0.80, *P* = 0.012). After accounting for changes in measured FC modularity without GSR, model-generated FC modularity change showed no clear additional association with either cognitive score (both *P* = 0.485).

## 4 Discussion

This exploratory study identifies differences in fitted network organization between tumour groups and relates postoperative changes in functional organization to cognitive change in meningiomas. Gliomas showed lower weighted GEC clustering than controls and meningiomas, accompanied by higher incoming participation. In meningiomas, changes in FC modularity without GSR were associated with changes in the attention/planning/ spatial-span score. In eligible glioma participants, lower FC between tumour regions and their counterparts provided anatomical context.

Weighted GEC clustering reflects both the presence of triangles in the fitted network and the weights of their connections. Higher incoming participation in gliomas indicates a more even distribution of incoming strength across the communities estimated for each participant (Fig. 2). Earlier work in this cohort reported higher structural participation in gliomas than in controls and examined model parameters describing global coupling and local inhibition [5]. Whole-brain models can estimate interregional connection weights by fitting functional observations [19]. In stroke, Idesis et al. [24] used fitted connectivity to examine altered network communication and showed that incorporating structural disconnection improved the reproduction of measured FC. Our analysis examines how fitted weights are organized across tumour groups, and checks whether the group difference holds under different structural and functional inputs.

Clustering remained lower in gliomas than in controls and meningiomas when each participant’s own SC was used as the GEC starting matrix. The common control SC alone therefore does not explain the finding. To assess the influence of the functional targets, we then kept each participant’s SC starting matrix fixed and replaced their FC and time-lagged covariance with control averages. Mean GEC clustering then fell in all three groups, most in controls, and became similar across groups. The groups therefore converged at a lower level, not at the level seen in controls fitted to their own targets. With control-average targets, all participants also share the same functional inputs, so the comparison removes between-participant variation as well. Together, these sensitivity analyses indicate that the clustering difference is not produced by the choice of starting matrix and arises from participants’ own functional targets.

At the regional level, FC between tumour regions and their counterparts was lower than for the same connections in controls (Fig. 3). This held for all ten eligible glioma participants, on average across each participant’s connections, both without and with GSR. Daniel et al. reported lower homotopic FC in normal-appearing cortex in patients with high-grade glioma than in controls [25]. Nenning et al. described distributed functional abnormalities involving both hemispheres in glioblastoma [26]. Our finding adds evidence of lower FC in connections that involve tumour regions; it does not show that lower FC is confined to those connections.

For the same tumour–counterpart pairs, standardized SC strength was lower at tumour regions than at their counterparts in nine of the ten patients, although the confidence interval for the mean SC strength difference included zero. Patients with more negative SC strength differences also tended to have lower global GEC clustering. This association offers a possible link between regional structural alterations and whole-brain fitted-network organization. The link remains tentative because the confidence interval for the correlation included zero and the Pearson analysis was inconclusive.

The longitudinal findings relate changes in functional network organization to cognitive change in meningiomas (Fig. 4). Correlations between FC modularity change and overall cognitive change were positive both without and with GSR, but remained inconclusive. An exploratory analysis of two subscores showed that FC modularity change without GSR was positively associated with change in the attention/planning/spatial-span score. This association remained after accounting for preoperative values of that cognitive score and FC modularity.

In a cross-sectional study of patients with resected WHO grade I meningiomas, van Nieuwenhuizen et al. reported exploratory associations between working memory and functional network organization measured with magnetoencephalography (MEG) [27]. Ellis et al. subsequently reported a positive association between FC modularity change and change in a composite neuropsychological score in a multivariable analysis of a heterogeneous surgical cohort assessed early after surgery [13]. Our study extends this question to changes over several months in meningiomas. Cognitive measures differed between the studies. We also estimated communities separately for each FC network, whereas Ellis et al. used predefined communities.

Modularity change in model-generated FC was also positively associated with change in the attention/planning/ spatial-span score. Because the networks were fitted to the measured functional targets, this shows that the model output is consistent with the measurements; it is not independent evidence. However, modularity change in model-generated FC showed no clear additional association with cognitive change after accounting for modularity change in measured FC without GSR.

Reporting FC without and with GSR allowed us to assess sensitivity to fluctuations shared across the cortex. Lower tumour–counterpart FC was present in both conditions. The association between FC modularity change and attention/planning/spatial-span change was weaker and inconclusive with GSR. In the baseline-adjusted analysis of that score, the estimate was similar in magnitude but less precise with GSR. Shared cortical fluctuations can contain both neural and non-neural contributions [14]. We therefore interpret the two conditions as complementary analyses of the same recordings. Differences in statistical support alone do not establish that the cognitive associations differ between conditions.

The small, heterogeneous cohort limits the precision and generalizability of the findings, particularly for the longitudinal analyses. Thirteen of the 14 meningiomas were WHO grade I, so the findings may not extend to higher-grade meningiomas. Differences in age, sex and acquisition TR also complicate interpretation of group contrasts. Covariate adjustment weakened the evidence for the whole-cortex glioma–control FC difference with GSR. For the comparison between fits to individual and control-average functional targets, statistical support for group differences depended on which covariates were included (Supplementary Table S1). These results make it difficult to attribute the observed group differences to tumour type alone. Tumour overlap was estimated using a common atlas and may not precisely reflect each participant’s anatomy. GEC weights are model estimates: they depend on the model, the starting matrix, the functional targets and the fitting settings, and are not direct measurements of anatomical connections. Associations between changes do not show that cognition improved after surgery, or that changes in FC modularity caused changes in cognition.

In summary, fitted networks in gliomas were less clustered than those of controls and meningiomas, a difference that held when each participant’s own SC was used for fitting. In meningiomas, changes in FC modularity after surgery accompanied changes in the attention/planning/spatial-span score. Lower FC between tumour regions and their counterparts adds anatomical context. Independent studies should now test whether the glioma pattern in fitted connectivity recurs across cohorts and whether changes in FC modularity consistently accompany cognitive change after surgery.

## Data availability

All data used in this study are publicly available from OpenNeuro: the BTC_preop dataset (ds001226, version 5.0.1) [15] and the BTC_postop dataset (ds002080, version 4.0.1) [16]. We used the participant tables and the derivatives supplied with the datasets: structural connectivity matrices, regional BOLD signals and tumour masks. The datasets and their acquisition are described in [5, 7, 17]. Upon acceptance, the analysis code, with instructions for obtaining the data and running the analyses, will be made available at github.com/ajunca/brain-tumor-connectivity.

## Acknowledgements

We thank Hannelore Aerts, Daniele Marinazzo and their colleagues at Ghent University for collecting and publicly sharing the BTC_preop and BTC_postop datasets [15, 16] analysed in this study, and the patients and control participants who made this research possible.

Large language model tools were used at various stages of the analysis and preparation of this manuscript. The authors reviewed all content generated by these tools and take full responsibility for it.

## Supplementary material

Supplementary material provides GEC group-comparison statistics and covariate-adjusted sensitivity analyses of whole-cortex FC and the GEC functional-target difference.

## References

[1] David N. Louis et al. “The 2016 World Health Organization Classification of Tumors of the Central Nervous System: a summary”. In: Acta Neuropathologica 131.6 (June 2016), pp. 803–820. ISSN: 0001-6322, 1432-0533. DOI: 10.1007/s00401-016-1545-1.

[2] Christian Ogasawara, Brandon D. Philbrick, and D. Cory Adamson. “Meningioma: A Review of Epidemiology, Pathology, Diagnosis, Treatment, and Future Directions”. In: Biomedicines 9.3 (Mar. 21, 2021), p. 319. ISSN: 2227-9059. DOI: 10.3390/biomedicines9030319.

[3] V.M. Stoecklein et al. “Perifocal Edema in Patients with Meningioma is Associated with Impaired Whole-Brain Connectivity as Detected by Resting-State fMRI”. en. In: American Journal of Neuroradiology 44.7 (July 2023), pp. 814–819. ISSN: 0195-6108, 1936-959X. DOI: 10.3174/ajnr.A7915.

[4] Erica Silvestri et al. “Widespread cortical functional disconnection in gliomas: an individual network mapping approach”. en. In: Brain Communications 4.2 (Apr. 2022), fcac082. ISSN: 2632-1297. DOI: 10.1093/braincomms/fcac082.

[5] Hannelore Aerts et al. “Modeling Brain Dynamics in Brain Tumor Patients Using the Virtual Brain”. In: eneuro 5.3 (May 2018), ENEURO.0083–18.2018. ISSN: 2373-2822. DOI: 10.1523/ENEURO.0083-18.2018.

[6] Albert Juncà et al. “Impact of meningioma and glioma on whole-brain dynamics”. en. In: Scientific Reports 16.1 (Jan. 2026), p. 5032. ISSN: 2045-2322. DOI: 10.1038/s41598-026-35140-1.

[7] Hannelore Aerts et al. “Modeling brain dynamics after tumor resection using The Virtual Brain”. In: NeuroIm-age 213 (June 2020), p. 116738. ISSN: 10538119. DOI: 10.1016/j.neuroimage.2020.116738.

[8] Milou Straathof et al. “A systematic review on the quantitative relationship between structural and functional network connectivity strength in mammalian brains”. en. In: Journal of Cerebral Blood Flow & Metabolism 39.2 (Feb. 2019), pp. 189–209. ISSN: 0271-678X, 1559-7016. DOI: 10.1177/0271678X18809547.

[9] Hualou Liang and Hongbin Wang. “Structure-Function Network Mapping and Its Assessment via Persistent Homology”. en. In: PLOS Computational Biology 13.1 (Jan. 2017). Ed. by Danielle S. Bassett, e1005325. ISSN: 1553-7358. DOI: 10.1371/journal.pcbi.1005325.

[10] Morten L. Kringelbach et al. “Toward naturalistic neuroscience: Mechanisms underlying the flattening of brain hierarchy in movie-watching compared to rest and task”. en. In: Science Advances 9.2 (Jan. 2023), eade6049. ISSN: 2375-2548. DOI: 10.1126/sciadv.ade6049.

[11] Mikail Rubinov and Olaf Sporns. “Complex network measures of brain connectivity: Uses and interpretations”. In: NeuroImage 52.3 (Sept. 2010), pp. 1059–1069. ISSN: 10538119. DOI: 10.1016/j.neuroimage.2009.10.003.

[12] Olaf Sporns. “Structure and function of complex brain networks”. In: Dialogues in Clinical Neuroscience 15.3 (Sept. 30, 2013), pp. 247–262. ISSN: 1958-5969. DOI: 10.31887/DCNS.2013.15.3/osporns.

[13] David G. Ellis et al. “Longitudinal changes in brain connectivity correlate with neuropsychological testing in brain tumor resection patients”. In: Frontiers in Neuroscience 19 (Mar. 2025), p. 1532433. ISSN: 1662-453X. DOI: 10.3389/fnins.2025.1532433.

[14] Kevin Murphy and Michael D. Fox. “Towards a consensus regarding global signal regression for resting state functional connectivity MRI”. en. In: NeuroImage 154 (July 2017), pp. 169–173. ISSN: 10538119. DOI: 10.1016/j.neuroimage.2016.11.052.

[15] H Aerts and D Marinazzo. BTC_preop. OpenNeuro, 2026. DOI: 10.18112/openneuro.ds001226.v5.0.1.

[16] Hannelore Aerts and Daniele Marinazzo. BTC_postop. OpenNeuro, 2026. DOI: 10.18112/openneuro.ds002080.v4.0.1.

[17] Hannelore Aerts et al. “Pre- and post-surgery brain tumor multimodal magnetic resonance imaging data optimized for large scale computational modelling”. In: Scientific Data 9.1 (Nov. 5, 2022), p. 676. ISSN: 2052-4463. DOI: 10.1038/s41597-022-01806-4.

[18] Rahul S. Desikan et al. “An automated labeling system for subdividing the human cerebral cortex on MRI scans into gyral based regions of interest”. In: NeuroImage 31.3 (July 2006), pp. 968–980. ISSN: 10538119. DOI: 10.1016/j.neuroimage.2006.01.021.

[19] Gustavo Deco et al. “Violations of the fluctuation-dissipation theorem reveal distinct nonequilibrium dynamics of brain states”. en. In: Physical Review E 108.6 (Dec. 2023), p. 064410. ISSN: 2470-0045, 2470-0053. DOI: 10.1103/PhysRevE.108.064410.

[20] Giorgio Fagiolo. “Clustering in complex directed networks”. en. In: Physical Review E 76.2 (Aug. 2007), p. 026107. ISSN: 1539-3755, 1550-2376. DOI: 10.1103/PhysRevE.76.026107.

[21] Vincent D Blondel et al. “Fast unfolding of communities in large networks”. In: Journal of Statistical Mechanics: Theory and Experiment 2008.10 (Oct. 2008), P10008. ISSN: 1742-5468. DOI: 10.1088/1742-5468/2008/10/P10008.

[22] Aric A. Hagberg, Daniel A. Schult, and Pieter J. Swart. “Exploring Network Structure, Dynamics, and Function using NetworkX”. In: Proceedings of the 7th Python in Science Conference (SciPy 2008). Pasadena, California, June 2008, pp. 11–15. DOI: 10.25080/TCWV9851.

[23] Pauli Virtanen et al. “SciPy 1.0: fundamental algorithms for scientific computing in Python”. en. In: Nature Methods 17.3 (Mar. 2020), pp. 261–272. ISSN: 1548-7091, 1548-7105. DOI: 10.1038/s41592-019-0686-2.

[24] Sebastian Idesis et al. “Inferring the dynamical effects of stroke lesions through whole-brain modeling”. In: NeuroImage: Clinical 36 (2022), p. 103233. ISSN: 22131582. DOI: 10.1016/j.nicl.2022.103233.

[25] Andy G S Daniel et al. “Homotopic functional connectivity disruptions in glioma patients are associated with tumor malignancy and overall survival”. en. In: Neuro-Oncology Advances 3.1 (Nov. 2021), vdab176. ISSN: 2632-2498. DOI: 10.1093/noajnl/vdab176.

[26] Karl-Heinz Nenning et al. “Distributed changes of the functional connectome in patients with glioblastoma”. In: Scientific Reports 10.1 (Oct. 27, 2020), p. 18312. ISSN: 2045-2322. DOI: 10.1038/s41598-020-74726-1.

[27] David Van Nieuwenhuizen et al. “Cognitive functioning and functional brain networks in postoperative WHO grade I meningioma patients”. en. In: Journal of Neuro-Oncology 140.3 (Dec. 2018), pp. 605–613. ISSN: 0167-594X, 1573-7373. DOI: 10.1007/s11060-018-2987-1.

