## supplemental tables for "Brain network organization and postoperative cognitive change in meningioma and glioma"

### Supplementary material

#### Covariate sensitivity analyses

For the seven comparisons in Table S1, we fitted separate regressions containing group, age and TR, then added sex to the same models. Age was centred within each comparison and scaled per ten years; TR was coded as 2.4 versus 2.1 s and sex as female versus male. All models included an intercept. Participants and outcomes were unchanged. Residual degrees of freedom were $n-4$ and $n-5$, respectively.

Whole-cortex mean absolute FC used ordinary least squares with HC3 standard errors and approximate two-sided $t$ tests and pointwise 95% intervals. The GEC outcome was clustering with individual functional targets minus clustering with control-average functional targets, both fitted from individual SC. Its group coefficient was assessed using 99,999 Freedman–Lane residual permutations within TR groups [1]. The reduced model contained all adjustment variables, including sex when applicable. The same permutations were used for both specifications; $P = (b+1)/100{,}000$, where $b$ counts absolute null $t$ statistics at least as large as observed. Fitted networks and shared inputs were held fixed.

BH correction covered the two FC comparisons within each GSR condition and the three GEC comparisons, separately for each specification. It did not cover selection among models. The small sample and common-slope assumptions limit interpretation; GEC permutation inference additionally assumes approximately homogeneous, exchangeable errors within TR groups. Its adjusted mean comparisons differ from the main rank tests.

Table S1: Group comparisons under two covariate-adjustment specifications.

| | Age + TR | | | Age + TR + sex | | |
|---|---|---|---|---|---|---|
| Comparison | Difference (95% CI) | Raw $P$ | BH $P$ | Difference (95% CI) | Raw $P$ | BH $P$ |
| **A. Whole-cortex FC without GSR** | | | | | | |
| Meningioma–control | −0.052 [−0.170, +0.065] | 0.367 | 0.444 | +0.011 [−0.075, +0.098] | 0.785 | 0.785 |
| Glioma–control | −0.053 [−0.195, +0.089] | 0.444 | 0.444 | −0.054 [−0.202, +0.095] | 0.457 | 0.785 |
| **B. Whole-cortex FC with GSR** | | | | | | |
| Meningioma–control | −0.012 [−0.063, +0.039] | 0.639 | 0.639 | −0.009 [−0.068, +0.050] | 0.747 | 0.747 |
| Glioma–control | −0.045 [−0.089, −0.001] | 0.045 | 0.090 | −0.045 [−0.095, +0.005] | 0.076 | 0.152 |
| **C. GEC functional-target difference in clustering** | | | | | | |
| Glioma–control | −0.030 | 0.035 | 0.063 | −0.030 | 0.029 | 0.043 |
| Meningioma–control | −0.018 | 0.154 | 0.154 | −0.001 | 0.929 | 0.929 |
| Meningioma–glioma | +0.025 | 0.042 | 0.063 | +0.032 | 0.021 | 0.043 |

Differences are adjusted regression coefficients, first group minus second. Controls: $n = 11$; meningiomas: $n = 14$; gliomas: $n = 11$. FC differences are in mean absolute correlation units; GEC differences are in clustering units. FC intervals are pointwise and are not BH-adjusted. Intervals were not estimated for the adjusted GEC coefficients. Raw $P$ denotes the two-sided test probability; BH $P$ its correction within each block and specification. These models assess sensitivity and do not establish tumour effects independent of demographic or acquisition differences.

Table S2: Group comparisons of GEC measures and the functional-target difference in clustering.

**A. GEC measures**

| Measure | Comparison | U | $r_{rb}$ | $P$ |
|---|---|---|---|---|
| Clustering | Glioma–control | 19 | −0.69 | 0.021 |
| | Meningioma–control | 55 | −0.29 | 0.239 |
| | Meningioma–glioma | 121 | +0.57 | 0.026 |
| Incoming participation | Glioma–control | 94 | +0.55 | 0.045 |
| | Meningioma–control | 72 | −0.06 | 0.805 |
| | Meningioma–glioma | 31 | −0.60 | 0.038 |
| Outgoing participation | Glioma–control | 92 | +0.52 | 0.063 |
| | Meningioma–control | 74 | −0.04 | 0.891 |
| | Meningioma–glioma | 34 | −0.56 | 0.060 |
| Global efficiency | Glioma–control | 25 | −0.59 | 0.065 |
| | Meningioma–control | 42 | −0.45 | 0.088 |
| | Meningioma–glioma | 98 | +0.27 | 0.262 |
| Clustering (individual SC) | Glioma–control | 19 | −0.69 | 0.021 |
| | Meningioma–control | 57 | −0.26 | 0.286 |
| | Meningioma–glioma | 117 | +0.52 | 0.046 |

**B. Functional-target difference in clustering**

| Comparison | Mean difference (95% CI) | U | $r_{rb}$ (95% CI) | $P$ |
|---|---|---|---|---|
| Glioma–control | −0.039 [−0.059, −0.018] | 20 | −0.67 [−0.95, −0.27] | 0.026 |
| Meningioma–control | −0.018 [−0.040, +0.004] | 51 | −0.34 [−0.74, +0.14] | 0.163 |
| Meningioma–glioma | +0.021 [+0.003, +0.039] | 118 | +0.53 [+0.12, +0.87] | 0.040 |

Controls: $n = 11$; meningiomas: $n = 14$; gliomas: $n = 11$. All tests are two-sided Mann–Whitney tests. U refers to the first named group; positive rank-biserial correlations ($r_{rb}$) indicate higher values in that group. All $P$-values are BH-adjusted across the three comparisons within each measure and fitting condition, and separately across the three functional-target comparisons. These tests do not adjust for age, sex or TR.

In A, networks were fitted to individual functional targets using common control SC, except for the final clustering block, which used individual SC. In B, each participant's score was clustering with individual targets minus clustering with control-average targets, both fitted from individual SC. Mean differences compare these scores between groups, first group minus second. Intervals are pointwise 95% percentile-bootstrap intervals from 20,000 participant resamples within groups, with fitted networks and shared inputs held fixed. Covariate sensitivity is reported in Table S1.

### References


[1] Anderson M. Winkler, Gerard R. Ridgway, Matthew A. Webster, Stephen M. Smith, and Thomas E. Nichols. "Permutation inference for the general linear model". In: *NeuroImage* 92 (2014), pp. 381–397. DOI: 10.1016/j.neuroimage.2014.01.060.
